# Activation mechanism and activity of globupain, a thermostable C11 protease from the Arctic Mid-Ocean Ridge hydrothermal system

**DOI:** 10.1101/2023.04.04.535519

**Authors:** Victoria Røyseth, Brianna M Hurysz, Anna Kaczorowska, Sebastian Dorawa, Anita-Elin Fedøy, Hasan Arsin, Mateus Serafim, Olesia Werbowy, Tadeusz Kaczorowski, Runar Stokke, Anthony J O’Donoghue, Ida Helene Steen

## Abstract

Deep-sea hydrothermal vent systems with prevailing extreme thermal conditions for life offer unique habitats to source heat tolearant enzymes with potential new enzymatic properties. Here, we present the novel C11 protease *globupain*, prospected from a metagenome-assembled genome of uncultivated *Archaeoglobales* sampled from the Soria Moria hydrothermal vent system located on the Arctic Mid- Ocean Ridges. By sequence comparisons against the MEROPS-MPRO database, globupain showed highest sequence identity to C11-like proteases present in human gut and intestinal bacteria,. Successful recombinant expression in *Escherichia coli* of the active zymogen and 13 mutant substitution variants allowed assesment of residues involved in maturation and activity of the enzyme. For activation, globupain required the addition of DTT and Ca²⁺. When activated, the 52 kDa proenzyme was processed at Lys_137_ and Lys_144_ into a 12 kDa light- and 32 kDa heavy chain heterodimer. A structurally conserved His_132_/Cys_185_ catalytic dyad was responsible for the proteolytic activity, and the enzyme demonstrated the ability to activate *in-trans*. Globupain exhibited caseinolytic activity and showed a strong preference for arginine in the P1 position, with Boc-QAR- aminomethylcoumarin (AMC) as the best substrate out of a total of 17 fluorogenic AMC substrates tested. Globupain was thermostable (T_m activated enzyme_ = 94.51 ± 0.09°C) with optimal activity at 75 °C and pH 7.1. By characterizing globupain, our knowledge of the catalytic properties and activation mechanisms of temperature tolerant marine C11 proteases have been expanded. The unique combination of features such as elevated thermostability, activity at relatively low pH values, and ability to operate under high reducing conditions makes globupain a potential intriguing candidate for use in diverse industrial and biotechnology sectors.

## 1. Introduction

Proteases catalyze the hydrolysis of peptide bonds in proteins and are important in industrial applications (Gimenes et al., 2021). They are used in food and leather processing, as additives to detergents, as pharmaceuticals, and in biorefineries (Barzkar et al., 2018; García-Moyano et al., 2021). Proteases are among the most widely used enzymes globally, accounting for over 60 percent of all enzyme sales (Ward, 2011). Temperature-tolerant proteases offer the possibility for industrial processing at high temperatures by improving reaction rates, enhancing nongaseous reactant solubility, and reducing contamination for mesophiles (Barzkar et al., 2018; Kumar et al., 2000). Deep-sea hydrothermal vents sustain microorganisms at high temperatures (Nunoura et al., 2008; Pikuta et al., 2007; Kuwabara et al., 2007), so they are an interesting starting point for discovering new thermostable proteases (Barzkar et al., 2018). Moreover, the increasing sequence diversity of encoded proteases revealed in hydrothermal vent microorganisms (Dombrowski et al., 2018; Cheng et al., 2021; Li et al., 2015) offers considerable potential for discovering new and novel proteases with optimized catalytic properties that may support future innovations.

Proteases are remarkably diverse in terms of activity and the nucleophilic residues that participate in hydrolysis (Rawlings and Bateman, 2019). Clostripain is a well-characterized endopeptidase originating from the bacterium *Clostridium histolyticum* (accession MER0000831) and is a member of the C11 enzyme family. Peptidases in this family are characterized by the presence of a catalytic cysteine-histidine dyad with a preference for hydrolyzing arginine and lysine bonds in the P1 position of peptide bonds (Barrett and Rawlings, 1996; Ogle & Tytell, 1953; Labrou and Rigden, 2004). Clostripain-like proteases are synthesized as inactive zymogens that have various requirements for activation (Kembhavi et al., 1991; McLuskey et al., 2016). Some require divalent cations such as Ca^2+^ and/or reducing agents such as dithiothreitol (DTT) for activation and catalysis. Variance in the number of cleavage sites for activation is also observed, and in some cases, an amino acid linker peptide is removed (Gilles et al., 1979; Dargatz et al., 1993). Nevertheless, the resulting active peptidase will comprise of a light and heavy chain making up a macromolecular active heterodimer. *In-trans* activation has been demonstrated in some, while others activate *in-cis*, reflecting the accessibility of cleavage sites to neighboring peptidase activity (Herrou et al., 2016; Roncase et al., 2019; González- Páez et al., 2019; Roncase et al., 2017).

This report presents a C11 protease called globupain, with ‘*globu*’ representing its unclassified *Archaeoglobus* species origin and ‘*pain*’ depicting it as a clostripain homolog. The type species, *Archaeoglobus fulgidus* (Stetter, 1988), of genus *Archaeoglobus*, was one of the first archaea to have its genome sequenced 25 years ago (Klenk et al., 1997). It has since served as a model thermophilic archaeon and provided important information about archaeal DNA replication (Maisnier-Patin et al., 2002), DNA repair (Birkeland et al., 2002; Knævelsrud et al., 2010), thermostable enzymes (Steen et al., 2001; Madern et al., 2001) and enzymes of biotechnological relevance (Isupov et al., 2019; Palombarini et al., 2020). With globupain, we have discovered a novel archaeal clostripain-like protease with a complex activation mechanism. Its unique catalytic properties and high thermal stability make globupain a promising candidate for industrial applications.

## 2 Materials and methods

### 2.1 Environmental sampling, DNA extraction, and sequencing

The Soria Moria vent field is part of the Jan Mayen vent fields (JMVFs), located at the southern part of the Mohns Ridge (Pedersen et al., 2005) in the Norwegian-Greenland Sea (71.2°N, 5.5°W). The end-member fluids of white smokers in the Soria Moria vent field have a pH of 4.1 and a concentration of hydrogen sulfide of 4.1 mmol kg^-1^ (Dahle et al., 2015). In June of 2011, an *in-situ* titanium incubator\ (Stokke et al., 2020) consisting of one chamber filled with 2 g of dried krill shells (Nofima, Bergen, Norway), mixed with grained flange rock material (Dahle et al., 2015), was deployed at ∼ 30-35 cm below seafloor (blsf) in sediments at 716 m depth. The temperature was measured to be ∼ 40°C and ∼ 70°C at 20 and 30 cm blsf, respectively, indicating diffuse hydrothermal venting. The sample was recovered in July 2012, and the incubated material was immediately snap-frozen in liquid nitrogen and stored at – 80°C. DNA was extracted with FastDNA^TM^ SPIN Kit for Soil (MP Biomedicals, CA, USA) and sequenced at the Norwegian Sequencing Center in Oslo, NSC (www.sequencing.uio.no).

### 2.2 Metagenomic assembly, binning, and annotation

For the primary metagenome, one plate of 454 GS FLX Titanium shotgun reads (average read length; 730 bp) was sequenced and assembled using the Newbler assembler v.2.8 (Roche, Basel, Switzerland) with a minimum identity of 96 % over a minimum of 35 bases. In total, 0.9 million (75%) of the 454 raw reads were assembled into contigs resulting in 7448 contigs > 500 bp and an N50 contig size of 10887 bp. Open reading frame (ORF) predictions were made using Prodigal v2.60 (Hyatt et al., 2010) and screened against MEROPS (Rawlings et al., 2014) (2015; Release 9.13). Putative signal peptides were identified by SignalP v4.1 (Petersen et al., 2011). For the secondary metagenome, Illumina NovaSeq 150 bp paired-end reads were filtered and assembled using fastp v0.23.2 and MEGAHIT v1.2.9, respectively. Of the 290 million filtered reads, 90.8% mapped to the assembly using the bwa-mem aligner v.0.7.17 (Md et al., 2019). Metagenome-assembled genomes (MAGs) were binned and refined using MetaWrap v1.3.2 (Uritskiy et al., 2018), which included the binning tools, MetaBat2 v2.12.1 (Kang et al., 2015; 2019), MaxBin2 v2.2.6 (Wu et al., 2016) and CONCOCT v1.0.0 (Alneberg et al., 2014). Contamination and completeness of the MAGs were assessed with CheckM v1.0.7 (Parks et al., 2015). Furthermore, the taxonomic classification of the globupain-associated MAG (INS_M23_B45) was performed using the GTDB toolkit v2.1.0 (Chaumeil et al., 2019; 2022) with the GTDB release 207_v2 (Parks et al., 2018; 2020;2022). ORF predictions were made with Prodigal v2.6.0 (Hyatt et al., 2010) as part of the annotation workflow designed by Dombrowski et al. 2020. Cross-referencing the cloned globupain from the primary metagenome with the assembly from the secondary metagenome identified an ORF sharing 100% identity over 481 amino acids and 4 additional amino acids at the C-terminus (CFVD) .

### 2.3 Sequence alignment and three-dimensional modeling

Sequence alignment was made using the ESPript 3.0 utility (Robert & Gouet, 2014). The amino acid sequences of distapain (MER0095672), clostripain (MER0095672), thetapain (MER0028004), and\ PmC11 (MER0199417) were retrieved from the MEROPS database (Rawlings et al., 2018). The alignment was based on NCBI BLAST+ sequence similarity search results using the blastp program (Madeira et al., 2022) with MEROPS-MPRO sequences.

The globupain sequence (FASTA) was submitted to AlphaFold (software version 2.1.1), available at NMRbox (Maciejewski et al., 2017; https://nmrbox.nmrhub.org/), for prediction of a three-dimensional (3D) protein structure with atomic accuracy. AlphaFold (Jumper et al., 2021) modeled globupain structure was downloaded (Varadi et al., 2022), and the model’s alignment and analysis were assessed with PyMOL software (v0.99c) (DeLano, 2006). Lastly, the modeled structure was compared to a previously published crystallized clostripain structure (PDB ID: 4YEC; Roncase et al., 2017) template for comparison purposes.

### 2.4 Gene synthesis

Based on primary metagenome data, the globupain gene (GenBank accession OQ718499) was synthesized by GenScript (GenScript, NJ, USA) and codon-optimized for *Escherichia coli* expression (**Supplementary Work Sheet 1**). The gene was cloned (cloning site *Nde*I and *Xho*I) into pET-21A by GenScript, omitting the predicted 21 amino acid signal peptide (SignalP v. 6.0, Teufel et al., 2022). The resulting signal-free protein was extended with Met at the N-terminus, whereas the C-terminus was extended with Thr and Arg before the 6xHIS-tag (TRHHHHHH). For identification of amino acids in the catalytic dyad and in maturation (**Figure 1**), targeted amino acid residues (**Table 1**) were substituted with Ala, and the respective coding genes were synthesized and cloned by GenScript as described for the wild-type (WT) globupain.

**Figure 1.**
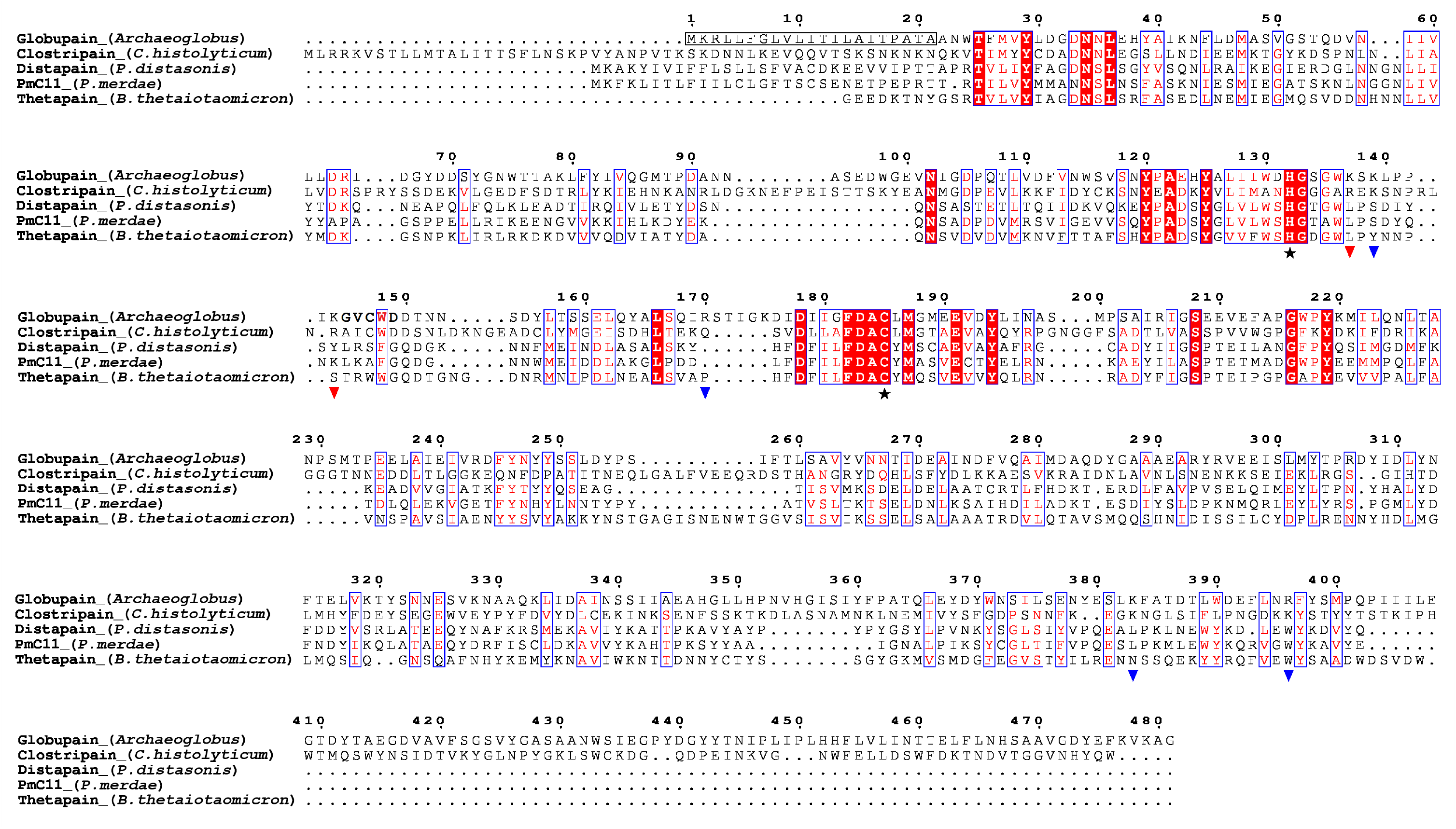
Sequence alignment of the C11 proteases globupain (*Archaeoglobales*), clostripain (*Clostridium histolyticum*), distapain (*Parabacteroides distasonis*), PmC11 (*P. merdae*), and thetapain (*Bacteroides thetaiotaomicron*). Symbols depict results from site-directed mutagenesis of the globupain coding sequence; ★, Cys/His catalytic dyad, ▼, sites showing resistance against cleavage when the amino acid was mutated into alanine, ▼, sites able to cleave when mutated into alanine. The detected N-terminal residues following activation are shown in bold.

**Table 1.** Overview of mutation variants of globupain and their targeted function

| Position in globupain | Putative function |
| --- | --- |
| Cys185Ala | catalytic dyad |
| His131Ala | catalytic dyad |
| Lys137Ala | self-activation |
| Lys139Ala | self-activation |
| Lys144Ala | self-activation |
| Lys137Ala/Lys139Ala | self-activation |
| Lys137Ala/Lys144Ala | self-activation |
| Lys139Ala/Lys144Ala | self-activation |
| Lys137Ala/Lys139Ala/Lys144Ala | self-activation |
| Arg170Ala | self-activation |
| Lys383Ala | cleavage C-terminal peptide |
| Arg396Ala | cleavage C-terminal peptide |
| Lys383Ala/ Arg396Ala | cleavage C-terminal peptide |

### 2.5 Protein production and purification

Expression plasmid of globupain and substitution variants were transformed into BL21-Gold (DE3) Chemically Competent *E. coli* cells (Agilent, TX, USA) using a heat-pulse manual supplied by the manufacturer. Cells were spread onto LB-agar plates supplemented with 100μg/mL ampicillin and incubated at 37°C overnight. Pre-cultures were inoculated by picking one single colony and incubating at 37°C in LB media containing 100μg/mL ampicillin with 190 rpm shaking overnight (Innova 44, New Brunswick Scientific, St Albans, United Kingdom). Expression cultures were inoculated with 5% of pre-culture in LB media with 100μg/mL ampicillin at 37°C and 190 rpm. At OD_600_ of 0.6, the temperature was set to 20°C, and the culture was equilibrated for 30 min. Heterologous expression was induced by IPTG brought to 0.1 mM IPTG, followed by overnight incubation (20°C). Cells were harvested by centrifugation at 5500 rpm for 15 min at 4°C. Pellets were stored at -20°C.

For purification of globupain and substitution variants, cells were resuspended in lysis buffer (50 mM HEPES, pH 7.5, 300 mM NaCl, 0.25 mg/ml lysozyme, 10% glycerol), placed on ice for 30 min, and lysed by ultra-sonication (5 times with 30% amplitude, in intervals of 20 sec on ice using the Vibra Cell with probe model CV188, Sonics and Materials INC, LT, USA). The lysate was clarified by centrifugation at 5500 rpm for 20 min at 4°C (Allegra^TM^ 21R Centrifuge, Beckman Coulter, CA, USA). The sample was then loaded into a HisTrap HP 5 mL column (Cytiva, Uppsala, Sweden) equilibrated with 20 mM HEPES, 500 mM NaCl, 25 mM imidazole, pH 7.5 with a flow rate of 1 mL/min. After elution with 20 mM HEPES, 500 mM NaCl, 500 mM imidazole, pH 7.5. fractions with the highest amount of enzyme were pooled and concentrated. The buffer was changed (20 mM HEPES, 150 mM NaCl, 0.1% CHAPS, pH 7.5) using Amicon® Ultra-15 centrifugal filter unit (Merck KGaA, Darmstadt, Germany) with a 30K molecular weight cut-off. Approximately 1 ml of the concentrated enzyme preparations were purified by gel filtration using a GE 16/600 Superdex 200 pg column (Cytiva, Uppsala, Sweden). Purified globupain and substitution variants were stored in 20 mM HEPES, 150 mM NaCl, 0.1% CHAPS, pH 7.5 at 4°C.

### 2.6 Maturation/activation

For activation of globupain and substitution variants (**Table 1**), a purified enzyme sample (< 5 mg/mL) was incubated at 75°C for up to 4.5 h in 20 mM tri-sodium citrate dihydrate, 150 mM NaCl, pH 5.5 (at room temperature) with 2.5 mM DTT and 1 mM CaCl_2_, respectively (activation buffer). To investigate if globupain could *in-tran*s activate, 10 μg of activated WT globupain was mixed with 10 μg of inactive C185A variant. The number and size of cleavage products were assessed by visualization of protein bands on 8-16% SurePAGE precast gels (GenScript) using MES SDS running buffer (GenScript) in a Bio-Rad Mini-PROTEAN Tetra Cell (BioRad, Hercules, CA, USA). For sample preparation, 4× lithium dodecyl sulfate (LDS) sample buffer (GenScript) with 2-mercaptoethanol was mixed with the protein sample, followed by denaturing at 95°C for 10 min. Gels were stained with InstantBlue^TM^ ultrafast protein stain (Abcam, Cambridge, United Kingdom), and the size of bands was indicated by broad multi-color pre-stained protein standard (GenScript). Edman sequencing was performed on a Shimadzu PPSQ-53A at the Iowa State University Protein Facility, USA.

### 2.7 Size-exclusion chromatography (SEC)

SEC analysis was performed using a Superdex 75 10/300 GL prepacked column connected to ÄKTA pure 25 chromatography system (GE Healthcare). The column was equilibrated with a 50 mM potassium phosphate buffer (pH 7.0), 150 mM NaCl and then loaded with a 500 µL sample of globupain protein (1 mg/mL). The flow rate of the run was adjusted to 0.5 mL/min, and the absorbance was measured at 280 nm (mAU, milli-absorbance units). For the experiment, the column was calibrated with proteins of known molecular weight: alcohol dehydrogenase (tetramer), 146,800; bovine serum albumin, 66,000; ovalbumin, 43,000; trypsin inhibitor, 22,000; and cytochrome C, 12,400 (Sigma-Aldrich, St. Louis, MO, USA). Dextran blue 2000 (Cytiva) was used to determine the column void volume.

### 2.8. Analytical ultracentrifugation (AUC)

Sedimentation velocity experiments were performed in a Beckman-Coulter ProteomeLab XL-I analytical ultracentrifuge (Indianapolis, IN, USA), equipped with AN 60Ti 4-hole rotor and 12 mm path length, double-sector charcoal-Epon cells, loaded with 400 μL of samples and 410 μL of buffer (50 mM potassium phosphate buffer pH 7.0, 150 mM NaCl, and 1 mM EDTA). The experiments were conducted at 20°C and 50,000 rpm, using continuous scan mode and radial spacing of 0.003 cm. Scans were collected in absorbance, in 4 min intervals at 280 nm. Data were analyzed using the “Continuous c(s) distribution” model of the SEDFIT program (Schuck, 2000), with a confidence level (F-ratio) specified to 0.6. Biophysical parameters of the buffer: density (1,01395 g/cm^3^), and viscosity (1,030 mPa s), were measured at 20°C using Anton Paar DMA 5000 density meter and Lovis 2000 ME viscometer. Protein partial specific volume (V-bars) was estimated at 0.7309 mL/g using SEDNTERP software (version 1.10, Informer Technologies Inc., Dallas, TX, USA). The results were plotted using GUSSI graphical program (Brautigam, 2015).

### 2.9 Thermal stability analysis

Thermostability of the inactive globupain and its activated form were assayed by nanoscale differential scanning fluorimetry (nanoDSF). Measurements were performed with Prometheus NT.48 instrument (NanoTemper Technologies, München, Germany) and PR.ThermControl software using standard grade capillaries. Before measurement, the capillaries were sealed with a sealing paste according to the manufacturer recommendations. The results were further analyzed with PR.StabilityAnalysis software. Thermostability of globupain zymogen at 0.4 mg/mL concentration and its activated form were assayed in 20 mM tri-sodium citrate (pH 5.5) buffer with 150 mM NaCl and 20 mM HEPES buffer pH 7.5 500 mM NaCl with 25 mM imidazole, respectively. Melting temperature of proteins was determined by thermal unfolding with a temperature gradient between 20-110°C at a ramp rate of 1°C/min. Thermal unfolding was measured by tryptophan and tyrosine fluorescence change at 330 and 350 nm emission wavelengths. All measurements were performed in triplicates.

### 2.10 Casein activity assays

The proteolytic activity of activated globupain and its substitution variants was assessed using the casein gelzan™ CM plate assay and the EnzChek™ Protease Assay Kit (Thermo Fisher Scientific, MA, USA), respectively. The gelzan™ CM plate assay was prepared by autoclaving 1.5% gelzan™ CM (Sigma-Aldrich) dissolved in 20 mM tri-sodium citrate (pH 5.5), 150 mM NaCl. The casein powder (Sigma-Aldrich) was dissolved in 20 mM tri-sodium citrate (pH 5.5), 150 mM NaCl, and autoclaved at 115°C for 10 min. Casein was added to the gelzan™ CM solution at a final concentration of 1.0%. The casein gelzan™ CM solution was poured into sterile glass Petri dishes and set to harden. Wells were made by punching holes in the plates using an inverted sterile 1 mL pipet tip. To test for proteolytic activity, 60 μL of activated globupain at 0.7-1.0 mg/mL was added to wells and incubated overnight at 75°C. Clearance zones would indicate caseinolytic activity.

When the proteolytic activity was assessed using the EnzChek™ Protease Assay Kit (Thermo Fisher Scientific), 20 mM tri-sodium citrate (pH 5.5) buffer with 150 mM NaCl was used to dilute the 1.0 mg/mL stock solution of BODIPY FL casein to 10 μg/mL. An aliquot of the activated enzyme (0.15 µg) was then added to the reaction mixture (100 μL of total volume) comprising 12.5 μL of 10 μg/mL BODIPY FL casein working solution and 77.5 μL of 20 mM tri-sodium citrate (pH 5.5) buffer with 150 mM NaCl, 10mM DTT, and 1 mM CaCl_2_. The caseinolytic activity was measured by running a time-resolved fluorescence read at 60°C, measuring fluorescence intensity every 20 s for 100 cycles. Fluorescence was measured with excitation wavelength 485 nm and emission wavelength 530 nm using an EnSpire™ 2300 Multilabel Reader (PerkinElmer, Turku, Finland).

### 2.11 Substrate screening

A total of 17 fluorogenic substrates containing a 7-amino-4-methylcoumarin (AMC) reporter group were screened for hydrolysis by Glopupain. These substrates consisted of Ac-VLTK-AMC, Ac- VLGK-AMC, Ac-VLVK-AMC, Ac-IK-AMC, Ac-YK-AMC, Ac-LK-AMC, Ac-LETK-AMC, Ac- IETK-AMC, Ac-AEIK-AMC, Ac-AIK-AMC, Boc-LRR-AMC (R&D Systems S-300), N-Benzoyl- FVR-AMC (Bachem I-1080), z-RR-AMC (Sigma C5429), Ac-RLR-AMC (AdipoGen AG-CP3-0013), Boc-QAR-AMC (Peptide International 3135-v), z-VVR-AMC (Peptide International 3211-v), Pyr-RTKR-AMC (Peptide International 3159-v) where the N-terminal blocking groups Ac, Boc, z, and Pyr, correspond to acetyl, tert-butyloxycarbonyl, benzyloxycarbonyl, and pyroglutamyl, respectively. Substrates containing K-AMC were synthesized by Dennis Wolan, The Scripps Research Institute, La Jolla, California and purified to >95%. All substrates were stored at -20°C as 10 mM stocks in DMSO.

Substrates were diluted to 100 μM in 20 mM tri-sodium citrate, 150 mM NaCl, 2.5 mM DTT, and 1mM CaCl_2_, pH 5.5, and mixed 1:1 with globupain such that the final concentration in the assay was of 2.9 μg/mL of enzyme and 50 μM substrate. Assays were performed in triplicate wells of a black 384-well plate (Thermo Fisher Scientific). Fluorescence was measured at 50°C over 1 h at excitation 360 nm emission 460 nm on a BioTek Synergy HTX Multimode Reader (BioTek, Agilent, Tx, USA). The reaction rate was calculated as the maximum velocity over 12 sequential readings, and means with standard errors were calculated. A Welch’s ANOVA and Brown Forsythe ANOVA were performed to calculate significances in GraphPad Prism 9.1.0. With Boc-QAR-AMC substrate, the Michaelis Menten kinetics was assessed at final concentrations ranging from 0 to 400 μM, and a Michaelis-Menten curve was fitted in GraphPad Prism 9.1.0 (**Supplementary Figure 1A**).

### 2.12 Determining the pH and temperature optimum

The pH optimum was determined using 5 nM globupain assayed against 50 μM Boc-QAR-AMC substrate in citrate phosphate buffer at various pH values. Samples were preincubated at 50°C for 10 min before fluorescence was measured. The optimum temperature for activity was assessed using Boc-QAR-AMC by incubating the enzyme and substrate at temperatures ranging from 25°C to 130°C in triplicate tubes. After 10 min, the enzyme was inactivated by mixing 1:5 in 8 M urea. Samples were plated on a black 384-well plate (Thermo Fisher Scientific), and the total fluorescence was measured at excitation 360 nm and emission 460 nm. The data reported is the average RFU for each temperature with standard error. Gaussian distribution was fitted in GraphPad Prism 9.1.0.

### 2.13 Data availability

The native C11 *globupain* protease has been submitted to GenBank under the accession number OQ718499. The sequence is archived under BioProject PRJNA296938 and derived from the primary metagenomic assembly, BioSample SAMN04111445. The reconstructed *Archaeoglobus* genome, INS_M23_B45, has been archived under the BioSample SAMN33944460 derived from the secondary metagenome, BioSample SAMN33925184. This Whole Genome Shotgun project has been deposited at DDBJ/ENA/GenBank under the accession JARQZL000000000. The version described in this paper is version JARQZL010000000.

## 3. Results

### 3.1 Metagenomic globupain discovery

By conducting deep-sea hydrothermal *in situ* enrichments emended with targeted biomass, we have previously shown the induced shifts in community structure towards higher fractions of heterotrophic microorganisms (Stokke et al., 2020). Furthermore, *in silico* screening from derived metagenomes has shown a high potential for discovering novel enzymes (Fredriksen et al., 2019; Stepnov et al., 2019; Vuoristo et al., 2019, Arntzen et al., 2021). In the current study, a novel protease named globupain was identified from an *in situ* enriched metagenome and targeted for expression and characterization. The selected gene encoded a C11 protease and originated from a metagenome-assembled genome (MAG) classified as an uncharacterized genome within the genus *Archaeoglobus* (INS_M23_B45; SAMN33944460 INS_M23_B45; SAMN33944460). The putative polypeptide comprised 481 AA with a 21-AA N-terminal signal peptide (**Figure 1**). The estimated molecular mass after signal peptide removal was 52.0 kDa, and the pI was 4.2, as determined by the ProtParam tool (Gasteiger et al., 2005). The highest sequence identity scores against the MEROPS-MPRO database (Rawlings et al., 2016) were of the human gut and intestinal C11 members; clostripain of 23.5% (*Clostridium histolyticum*), distapain of 27.3% (*Parabacteroides distasonis*), PmC11 of 24.2% (*P. merdae*) and thetapain of 26.9% (*Bacteroides thetaiotaomicron*). Sequence alignments (**Figure 1**) indicated a conserved catalytic His/Cys dyad in globupain at positions 132 and 185, respectively. Moreover, in the globupain model obtained with AlphaFold (**Figure 2A**) deep learning-based algorithm (Jumper et al., 2021), structural similarities were observed between the predicted structure and the available PmC11 crystal structure (**Figure 2B**;PDB ID: 4YEC). The active site residues in PmC11 (e.g., Asp_177_), including catalytic His_133_ and Cys_179_, were conserved between both structures (**Figure 2C**). The residues which did not overlay well with PmC11 include an area between the known heavy and light PmC11 chains (**Figure 2D**) and a long C-terminal region (**Figure 2E**).

**Figure 2.**
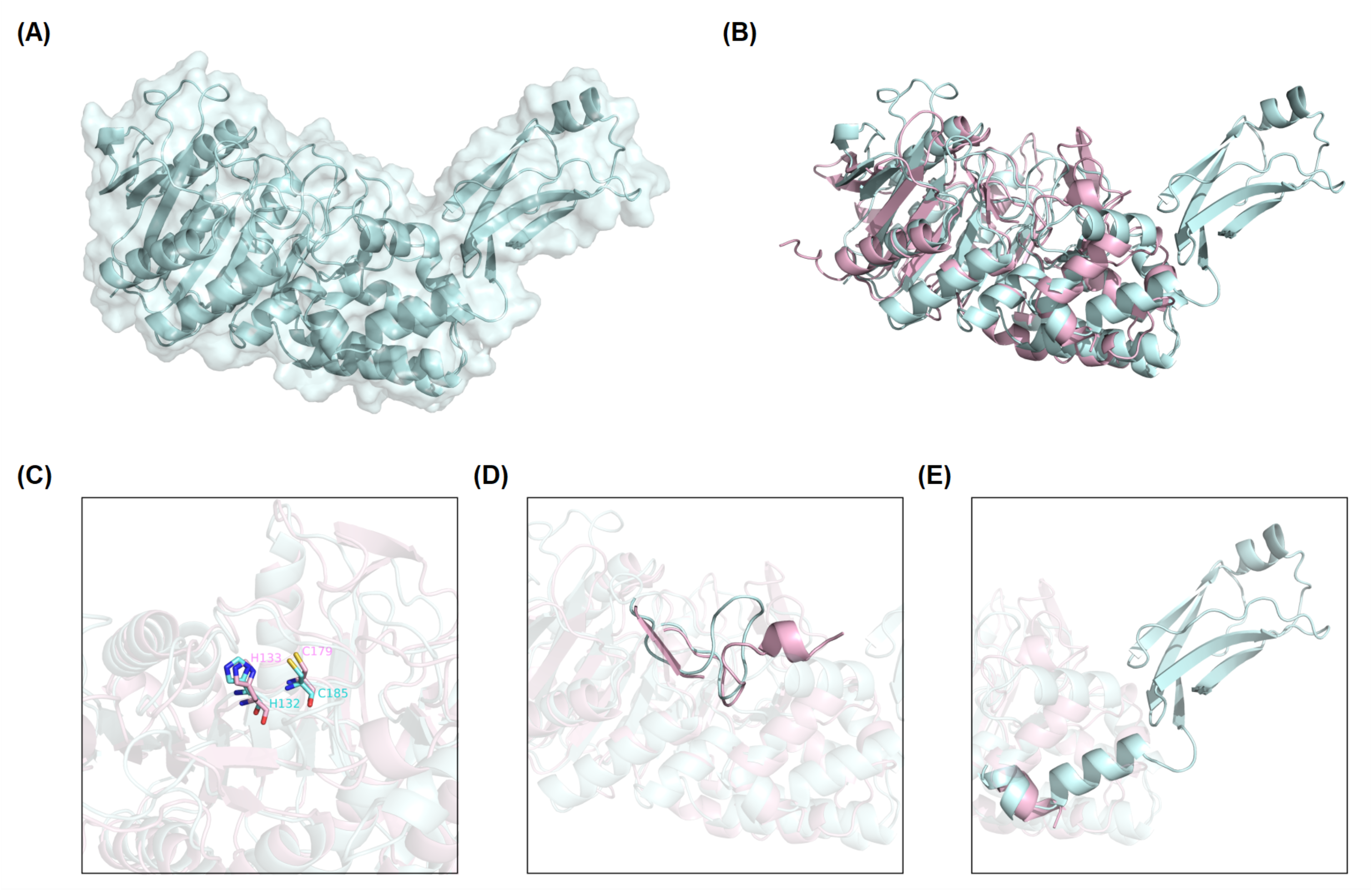
Globupain modeled structure predicted with AlphaFold in comparison to PmC11. (A) AlphaFold predicted structure is represented by cartoon with transparent surface (palecyan) (B) Alphafold of globupain compared to crystallized PmC11 (PDB ID: 4YEC) structure (lightpink) in a 3D alignment showing their structures similarities. (C) Active site residues (e.g., His, Asp and Cys) are conserved among the two aligned structures. (D) Light and heavy chain cleavage region is depicted for the modeled globupain superimposed to clostripain’s structure, as well as the (E) likely C-terminal cleavage region. Images were generated with PyMOL (v0.99c).

### 3.2 Globupain activation

Globupain and substitution variants were expressed as soluble proteins in *E. coli* BL21-Gold (DE3), with almost 100% of the total recombinant protein being in a soluble form (**Supplementary Figure 2, 3**). The wild-type enzyme was produced as an inactive zymogen. However, incubation at 75°C for 4.5 h (**Supplementary Figure 4**) in the activation buffer resulted in an active form of the C11 globupain. SDS-PAGE imaging showed that the 52 kDa zymogen was cleaved into a 32 kDa heavy chain and a 12 kDa light chain (**Figure 3A, B**), forming a heterodimer stabilized by noncovalent bonding. Oligomeric structure analysis performed by size-exclusion chromatography and analytical ultracentrifugation revealed that globupain in zymogen form exists in solution as a homodimer (**Supplementary Figure 5, 6**). While the zymogen is inactive, the enzyme, after activation, can hydrolyze casein (**Figure 3E, F**). Globupain activation and its enzymatic activity, as in the case of other C11 proteases (Labrou and Rigden, 2004), depends on the presence of His/Cys catalytic dyad in the primary protein sequence (H_132_/C_185_). H_132_ of the light chain is responsible for deprotonating the neighboring C_185_ in the heavy chain, which then promotes its nucleophilic attack on the substrate. For globupain, we found that substitution variant C_185_A generated by site-directed mutagenesis cannot be activated (**Figure 3C**). Also, no caseinolytic activity was observed for either H_132_A or C_185_A variants (**Figure 3E**).

**Figure 3.**
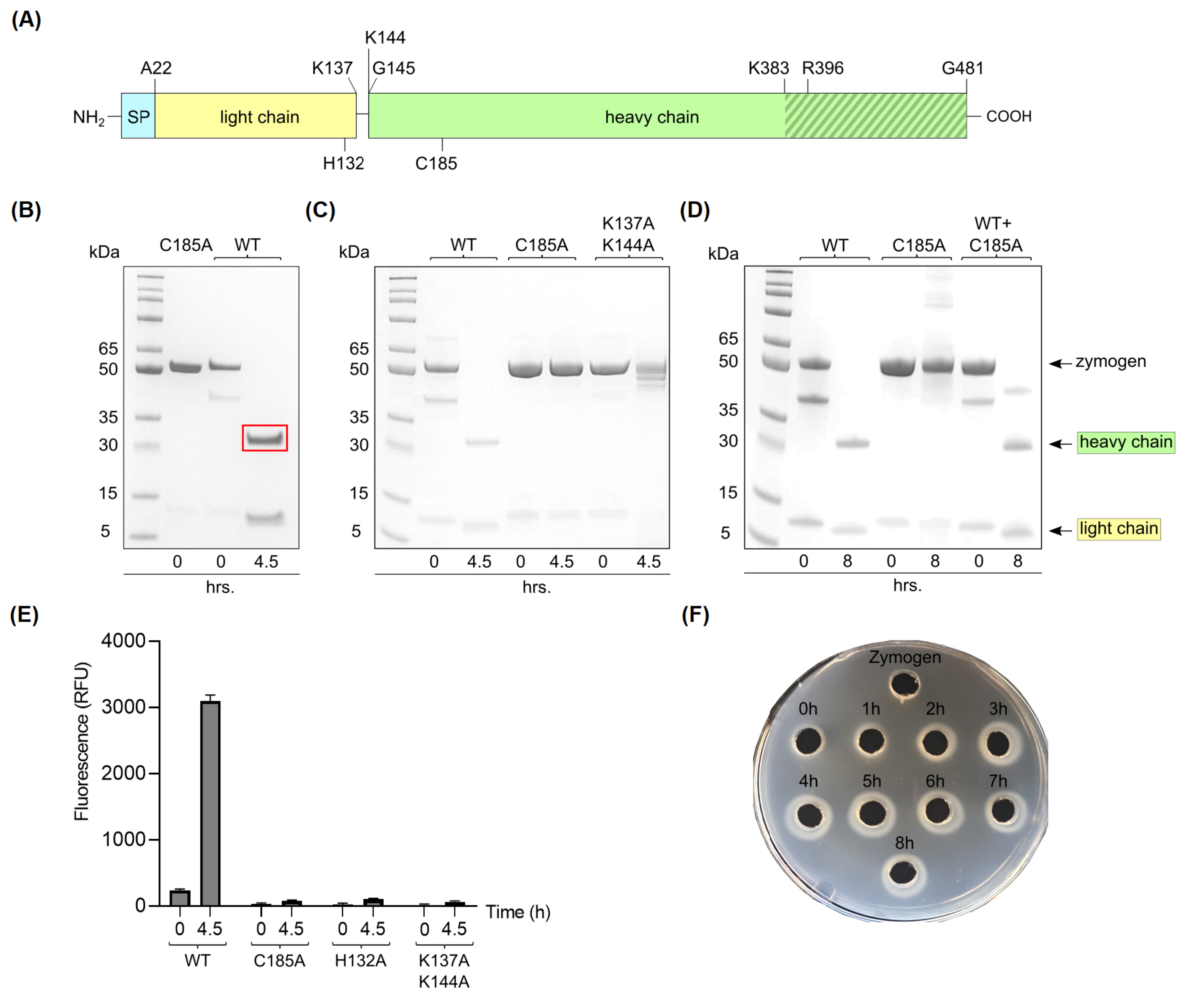
Activation of globupain. (A) A schematic representation of the primary structure of globupain. Globupain was overproduced without the N-terminal 21 amino acid signal peptide (SP). The light chain (yellow) and the heavy chain (green) of the active heterodimer result from zymogen activation. The cleavage sites at K_137_ and K_144_ are shown. H_132_ and C_185_ of the catalytic dyad are indicated on the light- and heavy chain, respectively. K_383_ and R_396_ located C-terminally on the heavy chain (a grey area), were not confirmed to be involved in zymogen activation. (B) SDS-PAGE gel presentation of inactive C185A and wild-type (WT) of 52 kDa, respectively, and activation of WT into a 32 kDa heavy- and 12 kDa light chain when incubated at 75°C for 4.5 hrs, The region within the red rectangle was excised for N-terminal sequencing. (C) SDS-PAGE gel analysis shows that the C185A variant and K_137_A/K_144_A variant cannot process into a heterodimer. (D) SDS-PAGE image of WT, C_185_A and WT+C_185_A incubated at 75°C at 0h and 8h in activation shows that the enzyme is able to *in- trans* activate. (E) When activated for 4.5h at 75°C in activation buffer, globupain can cleave casein whereas mutation variants H_132_A, C_185_A and K_137_A/K_144_A showed no increase in fluorescence (RFU) when assayed with EnzChek™ Protease Assay Kit at 60°C. (F) Casein-gelzan plate showing globupain zymogen and clearance zones when activated at 75°C for 0-8h.

Edman sequencing on a Shimadzu PPSQ-53A at the Iowa State University Protein Facility of the heavy chain revealed that the N-terminus consisted of G_145_VCWD; hence cleavage (*) occurred between K_144_ and G_145_ within the sequence LPPIK*GVCWD (**Figure 1**). To further evaluate globupain autoprocessing at this cleavage site, a K_144_A variant was constructed. Notably, processing of the zymogen into this variant’s heavy and light chain still occurred with similar size of cleaved products as WT globupain, as visualized by SDS-PAGE (**Supplementary Figure 7**). To further assess if this result could be explained by cleavage after nearby Lys residues, 7 new variants were synthesized (**Table 1, Supplementary Figure 7**). Only the double (K_137_A/K_144_A) and triple (K_137_A/K_139_A/K_144_A) mutants failed to activate into the processed form (**Figure 3C**) and remained catalytically inactive (**Figure 3E, Supplementary Figure 7**), which altogether suggests that globupain can self-activate by cleavage after both Lys_137_ and Lys_144_ (**Figure 1**), respectively.

When mixing the WT zymogen with inactive C_185_A and performing the standard activation protocol, both the WT and C_185_A proteins were processed into the light and heavy chain (**Figure 3D**). This finding demonstrates that globupain can activate *in trans* and indicates that the sites for activation are exposed for cleavage by nearby protease. Interestingly, the activation sites results in the removal of the unique region that poorly overlays with PmC11 seen in **Figure 2D**.

The combined molecular mass of the heavy and light chains of activated globupain was determined to be 44 kDa, which, when compared to the 52.0 kDa zymogen (**Figure 3A, B**), indicates additional autoprocessing occurs during activation. This discrepancy in molecular weight points to a likely cleavage in the C-terminal region, which the model supports (**Figure 2E**). Activated globupain (**Supplementary Figure 8**) failed to bind to the Ni²^+^ affinity column, and the C-terminal His-tag of globupain was not detected on either light or heavy chain (**Supplementary Figure 9**) altogether, revealing that a C-terminal fragment that contains the His-tag was removed during autoprocessing. Manual inspection of the primary sequence suggested that K_383_ and R_396_ may be the putative cleavage sites. However, each of the enzyme variants K_383_A, R_396_A, and K383A/R_396_A were still processed into the active form, and their C-terminal portion was removed (**Supplementary Figure 10**), indicating that K_383_ and R_396_ were not the sites for C-terminal processing.

### 3.3 Substrate specificity (AMC) determination

To quantify globupain activity in a microwell plate assay, the enzyme was incubated with three substrates that we previously developed for another clostripain-like C11 family member known as PmC11 (Roncase et al., 2017). This enzyme was encoded in the *Parabacterioides merdae* genome. The substrates consisted of tetrapeptides (VLXK) with an N-terminal acetyl group (Ac) and a C- terminal AMC reporter group. These substrates were chosen as the P1 residue corresponds to the N- terminal auto-activation site of globupain. For PmC11, Ac-VLTK-AMC was most efficiently cleaved, followed by Ac-VLGK-AMC. However, the substitution of the P2 residue for a hydrophobic Val side chain ablated substrate turnover by PmC11. We tested all three substrates against globupain and found them to be cleaved at a similar rate (**Figure 4A**). This finding revealed that globupain has broader substrate specificity to PmC11 at the P2 position. Subsequently, 7 other substrates with Lys at P1 available in the lab were tested (**Figure 4B**). Globupain was able to cleave all substrates and cleaved AIK-AMC with highest efficiency. Interestingly, globupain cleaved each of the 7 new substrates more efficiently than the set of three initial substrates, which were based on the optimal substrate of PmC11 from *P. merdae*, indicating a distinct specificity to PmC11. The most statistically significant cleavage differences (p < 0.01) amongst the new 7 substrates occurred between Ac-AIK-AMC (the most efficient) and Ac-YK-AMC (least efficient). Activity with Ac-AIK-AMC was also statistically significantly increased (p < 0.01) compared to Ac-AEIK-AMC. We hypothesized that the broad enzymatic activity of globupain for degrading casein may be partially due to cleavage following a structurally similar amino acid, arginine. Therefore, we examined fluorogenic substrates with Arg as the P1 residue (**Figure 4C**). This screen of 7 additional substrates revealed that Globupain cleaved substrates with even higher efficiency than the previous best with Lys at P1. However, some substrates, such as z-RR-AMC and Pyr-RTKR-AMC, showed minimal cleavage, indicating globupain may favor non-polar amino acids at the P2 position. Out of the 17 AMC substrates that were tested in this study, it was clear that globupain had a strong preference for R in the P1 position, with Boc-QAR-AMC as the best substrate.

**Figure 4.**
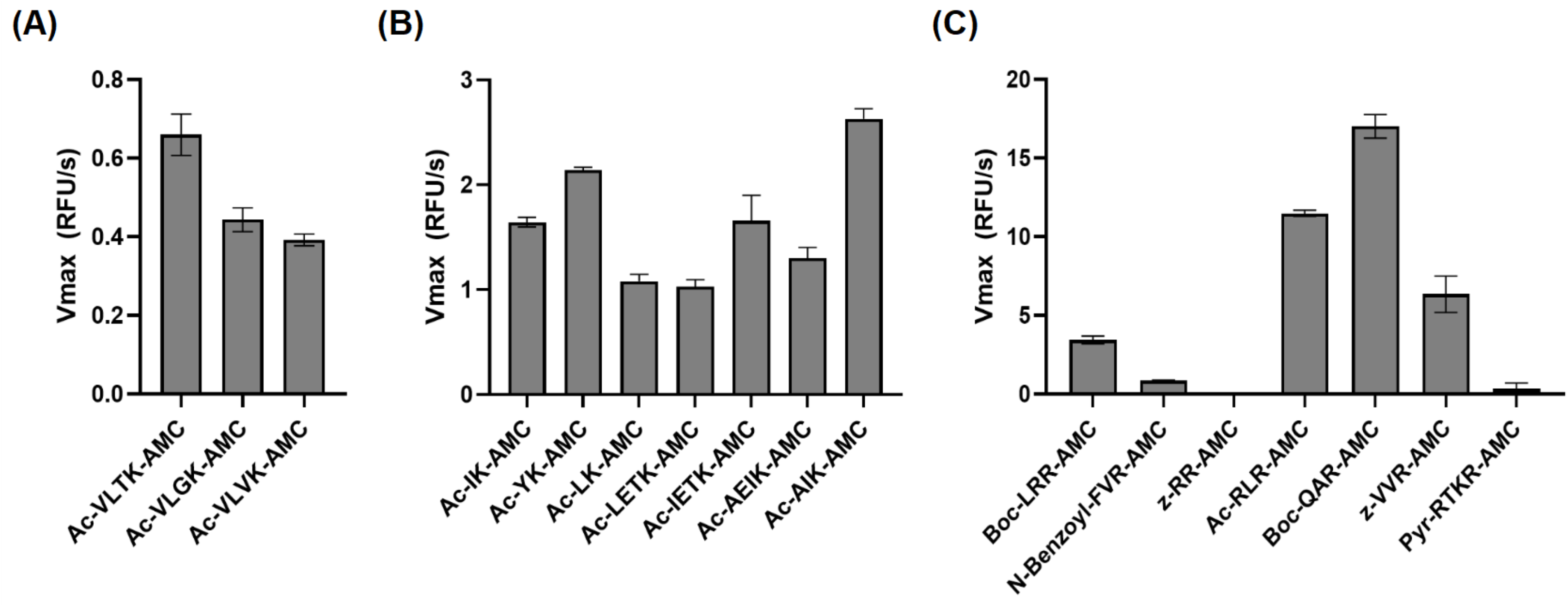
Substrate utilization by globupain. Globupain was assayed against 50uM of each fluorescent substrate at 50°C, fluorescence was measured, and the rate of enzyme cleavage, V_max_, for each substrate is reported. (A) Globupain was assayed against the substrates inititally designed for PmC11. (B) Globupain was assayed against 7 additional substrates with Lys at P1. (C) Globupain was assayed against 7 additional substrates with Arg at P1.

### 3.4 Thermal stability and optimal temperature and pH

Using the Boc-QAR-AMC substrate, the temperature optimum of globupain was determined to be 75.4 ± 0.56°C and remained 90% active at 60 and 90°C (**Figure 5A**). Thermal stability of inactive globupain as characterized by melting temperature (T_m_), which indicates the point at which half the protein is unfolded was 84.59 ± 0.21°C. The activated heterodimer’s melting temperature was 94.51 ± 0.09°C (**Figure 5B**). Finally, we evaluated the optimum pH of globupain using the Boc-QAR-AMC substrate. The optimum pH for catalytic activity was calculated to be pH 7.1 (**Figure 6A**). While the optimum pH is higher than the pH that we used for activation, the enzyme was shown to be more stable against autolysis at pH 5.5 than 7.1 (**Figure 6B,C,D**) which supported our use of pH 5.5 buffers for the biochemical characterization of globupain.

**Figure 5.**
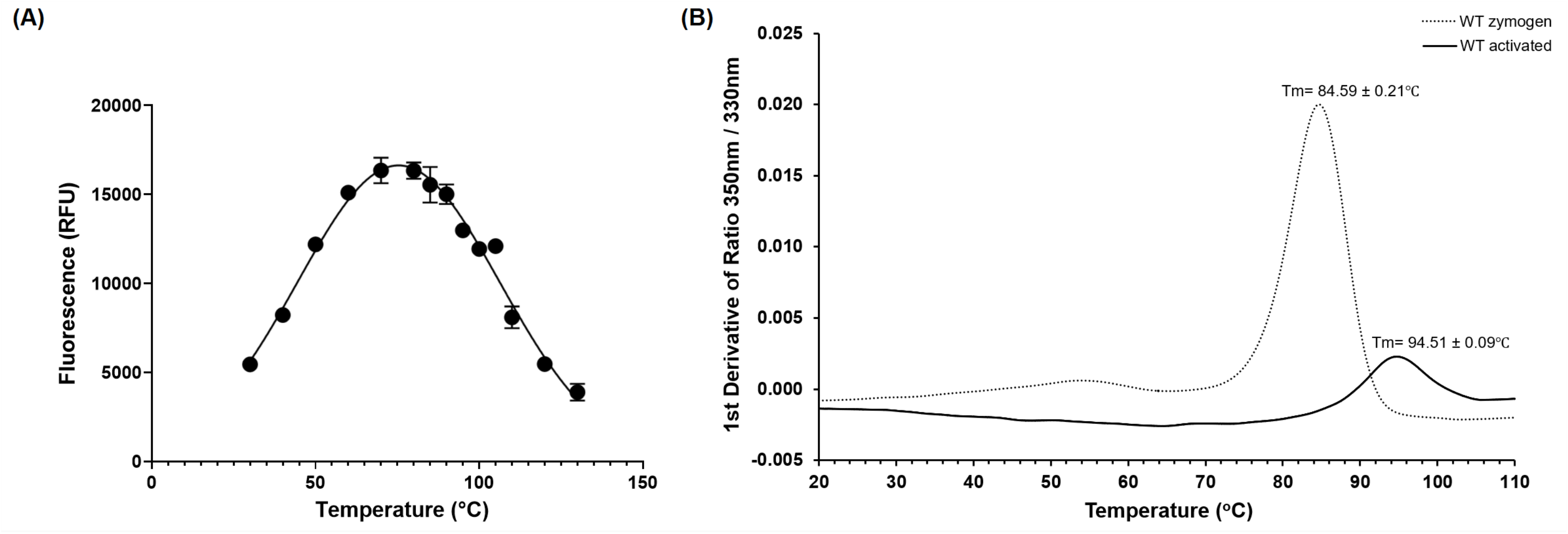
Themoactivity and thermostability of WT globupain. (A) Optimal temperature of globupain activity determined by incubating globupain with the lead substrate, Boc-QAR-AMC, at various temperatures and inactivating with urea before measuring fluorescence which correlated to substrate cleavage. (B) Thermogram of zymogen and activated form of globupain. Y-axis represents the first derivative of fluorescence intensity ratio 350 nm/330 measured by nanoDSF. Tm values are the mean and standard deviation from 3 replicate measurements.

**Figure 6.**
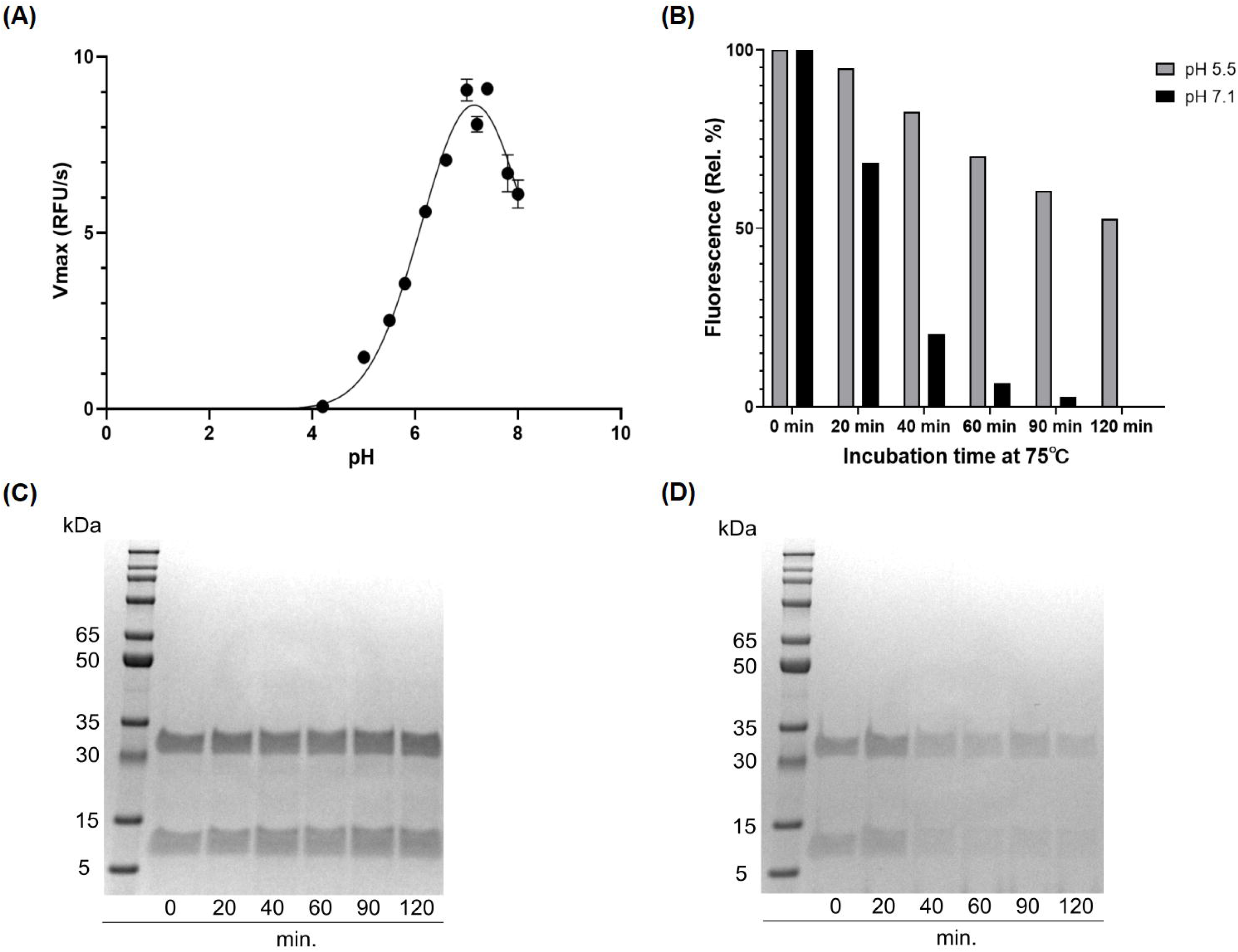
pH effect of activity and autolysis of globupain. (A) pH-optimum resolved by assaying with the substrate Boc-QAR-AMC in bufferes ranging from pH 2 to pH 8. Enzyme activity is shown as V_max_ for the different pH-values. (B) Time-dependent loss of enzyme activity (0.1 mg/mL) at pH 5.5. and 7.1, respectively. Enzyme activity was measured at 60°C with the with EnzChek™ Protease Assay Kit. (C) SDS-PAGE gel presentation reveals intact globupian after incubation at pH 5.5 wherease at pH 7.1 (D), autolysis is observed, explaining the loss of activity in (B).

## 4. Discussion

In this study, we characterized the novel cysteine protease, globupain belonging to the C11 enzyme family. Globupain was prospected from metagenomic data assigned to an unclassified *Archaeoglobus* species from the Arctic Mid-Ocean Ridge vent fields. The enzyme was highly soluble, expressing at relatively high concentrations in *E. coli*. Two protein bands (52 kDa and 44 kDa) with intact C-terminal His-tag were visualized on SDS-PAGE gels after protein purification. The zymogen of globupain is processed in the N-terminal region at Lys_137_ and Lys_144_ to yield a heavy and light chain. Similar to clostripain from *C. histolyticum* (Kembhavi et al., 1991), globupain requires calcium and a reducing environment for activation (**Supplementary Figure 4**). This condition contrasts the C11 protease from *P. merdae* PmC11, which activates independently of calcium (McLuskey et al., 2016). When activated, the globupain enzyme cleaves off a C-terminal peptide. The exact position for this remains unknown. This kind of autoprocessing is not uncommon for C11 proteases; for example, activation of clostripain starts with a 23 amino acid pro-peptide removal (Dargatz et al., 1993). Globupain has two cut sites at K_137_ and K_144_, leading to the removal of a 7-amino acid linker sequence and the formation of a heterodimer consisting of a heavy and light chain. For clostripain, a linker peptide is removed by cleavage at two Arg sites (Gilles et al., 1979; Dargatz et al., 1993). When activated, globupain showed the ability to *in-trans* activate and implied that the cut sites (K_137_ and K_144_) are exposed to proteolytic cleavage by neighboring proteases. This kind of activation is known to occur for several C11 enzymes such as thetapain (Roncase et al., 2019), fragipain (Herrou et al., 2016), and distapain (González-Páez et al., 2019) and contrasts PmC11 which activates only *in-cis* (Roncase et al., 2017). Globupain showed maximum activity at pH 7.1. This value is in the same range as known pH optima of PmC11 (pH 8.0), clostripain (pH 7.4-7.8), and thetapain (pH 7.4), respectively (Roncase et al., 2017; Ogle & Tytell, 1953; Mitchell & Harrington, 1968; Roncase et al., 2019). However, globupain showed an optimum temperature of 75°C and matures into a heat tolerant enzyme, which allows it to function in its thermal environment (Dahle et al., 2015). The oberserved thermal properties are in line with the growth characteristics of cultivated species within the genus *Archaeoglobus* (Mori et al., 2008; Steinsbu et al., 2010; Huber et al., 1997; Slobodkina et al., 2021; Burggraf et al., 1990; Stetter, 1988) and enzymes characterized previously (Steen et al., 2001). Moreover, in comparison to well-characterized industrially relevant, marine thermostable proteases, the thermal tolerance of globupain is superior to proteases sourced from marine *Bacillus* species and in the same range as of proteases from (hyper)thermophilic archaea (Barzkar et al., 2018).

Active clostripain-like proteases have been identified in marine sediment archaea (Lloyd et al., 2013). However, the highest sequence similarity scores of globupain using the MEROPS-MPRO database (Rawlings et al., 2016) were C11 proteases that originate from bacteria such as *C. histolyticum*, *P. distasonis*, *P. merdae, B. thetaiotaomicron* that have been found in the human intestinal microbiota (Salyers, 1984; Johnson et al., 1986; Franks et al., 1998). Some of these bacteria have been reported to cause disease and/or affect human health and have been studied to a greater extent (Salyers, 1984; Ezeji et al., 2021; McLuskey et al., 2016; Roncase et al., 2017; Roncase et al., 2019; Parracho et al., 2005). This finding highlights the significance of acquiring greater knowledge of marine C11 proteases. Notably, all C11 proteases, including globupain, show a conserved His/Cys catalytic dyad by sequence alignment. Moreover, the catalytic residues were also conserved in the globupain model obtained with AlphaFold (Jumper et al., 2021). Finally, it was shown experimentally using site-directed mutagenesis that in globupain, H_132_ and C_185_ were critical for activation and activity. When assayed against several AMC substrates, the enzyme showed a clear preference for the substrate Boc-Gln-Ala-Arg-AMC. Preference for hydrolyzing Arg bonds in the P1 position is a known trait for C11 members (Ogle & Tytell, 1953; Labrou & Rigden, 2004). It showed much lower activity against the Ac-Val-Leu-Thr-Lys-AMC substrate, which both PmC11 and thetapain hydrolyze efficiently (Roncase et al., 2017; Roncase et al., 2019). This observation indicates that the substrate specificity may vary substantially between different C11 proteases despite having sequence and structural similarities around the active site. In conclusion, the revealed temperature tolerance and catalytic properties of globupain render it as a promising protease in diverse industrial and biotechnology sectors. Further studies focused on in-depth knowledge of the substrate specificity (O’Donoghue et al., 2012, Rohweder et al., 2022), effects of protease inhibitors and resistance to organic solvents and chemical denaturants may provide a deeper understanding of the applicability of globupain.

## Author Contributions

VR, AJO, and IHS conceived the study. VR, BMH, and IHS wrote the manuscript. VR, BMH, AK, SD, AEF, HA, MS, OW, TK, and RS performed the experiments. All authors reviewed and edited the manuscript.

## Acknowledgments

We thank the cruise leader, professor Rolf-Birger Pedersen and the crew of G. O. SARS for their assistance during sampling campaigns 2011–2012 and Dennis Wolan for gifting us with several fluorescent substrates. The authors would also like to thank OpenEye Scientific for the academic licenses.

