## Supplementary data for "Activation mechanism and activity of globupain, a thermostable C11 protease from the Arctic Mid-Ocean Ridge hydrothermal system"

**Supplementary Work Sheet 1.** FASTA file of codon optimized globupain DNA sequence.

>Globupain_codon_optimized

ATGGCGAACTGGACCTTCATGGTGTACCTGGACGGCGATAACAACCTGGAGCACTATGC
GATCAAGAACTTTCTGGACATGGCGAGCGTTGGTAGCACCCAGGATGTGAACATCATTG
TTCTGCTGGACCGTATCGATGGTTACGACGATAGCTATGGCAACTGGACCACCGCGAAA
CTGTTCTACATTGTGCAGGGCATGACCCCGGACGCGAACAACGCGAGCGAGGATTGGGG
TGAAGTGAACATCGGCGACCCGCAAACCCTGGTGGATTTTGTTAACTGGAGCGTTAGCA
ACTACCCGGCGGAGCACTATGCGCTGATCATTTGGGACCACGGTAGCGGCTGGAAGAGC
AAACTGCCGCCGATCAAGGGTGTTTGCTGGGACGATACCAACAACAGCGATTACCTGAC
CAGCAGCGAACTGCAGTATGCGCTGAGCCAAATTCGTAGCACCATCGGTAAAGACATTG
ATATCATTGGCTTCGACGCGTGCCTGATGGGTATGGAGGAAGTGGATTACCTGATCAAC
GCGAGCATGCCGAGCGCGATTCGTATCGGCAGCGAGGAAGTTGAGTTTGCGCCGGGTTG
GCCGTATAAGATGATTCTGCAAAACCTGACCGCGAACCCGAGCATGACCCCGGAGGAAC
TGGCGATTGAAATCGTGCGTGACTTCTACAACTACTATAGCAGCCTGGATTATCCGAGC
ATCTTTACCCTGAGCGCGGTGTACGTTAACAACACCATCGACGAGGCGATTAACGATTT
CGTGCAGGCGATTATGGACGCGCAAGATTACGGTGCGGCGGCGGAAGCGCGTTATCGTG
TTGAGGAAATCAGCCTGATGTACACCCCGCGTGACTACATTGATCTGTATAACTTTACCG
AGCTGGTGAAGACCTATAGCAACAACGAAAGCGTTAAGAACGCGGCGCAGAAACTGAT
TGACGCGATCAACAGCAGCATCATTGCGGAGGCGCATGGTCTGCTGCACCCGAACGTTC
ACGGTATTAGCATCTACTTCCCGGCGACCCAACTGGAATACGATTATTGGAACAGCATC
CTGAGCGAGAACTATGAAAGCCTGAAATTTGCGACCGACACCCTGTGGGATGAGTTCCT
GAACCGTTTTTACAGCATGCCGCAGCCGATCATTATCCTGGAGGGCACCGACTATACCG
CGGAAGGTGATGTGGCGGTTTTCAGCGGTAGCGTGTATGGTGCGAGCGCGGCGAACTGG
AGCATCGAAGGTCCGTATGACGGCTACTATACCAACATTCCGCTGATCCCGCTGCACCA
CTTCCTGGTTCTGATTAACACCACCGAGCTGTTTCTGAACCACAGCGCGGCGGTGGGCG
ACTACGAATTTAAGGTTAAAGCGGGTCTCGAG

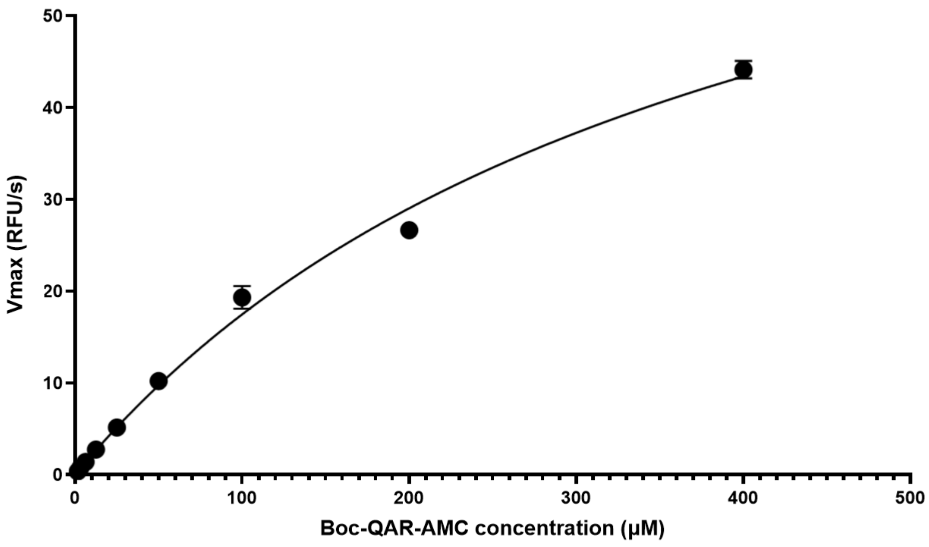


**Supplementary Figure 1.** Kinetic properties of WT-globupain. (A) Michaelis-Menten curve generated for globupain by varying lead substrate, Boc-QAR-AMC, concentration and measuring globupain enzyme activity via fluorescence which was then reported as $V_{max}$.

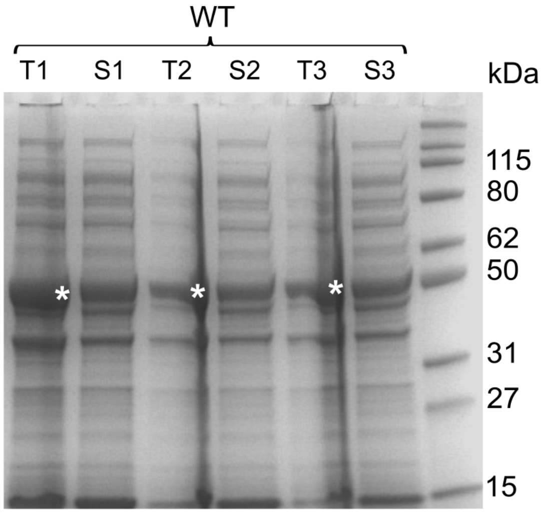


**Supplementary Figure 2.** Image of SDS-PAGE gel showing total (T) WT globupain expressed and the corresponding soluble protein (S). The figure shows triplicates (1-3). White asterisks indicate estimated protein size. To the far right shows the protein marker (Elite Pre-stained Protein Ladder, ProteinArk) with indicated molecular weight in kDa.

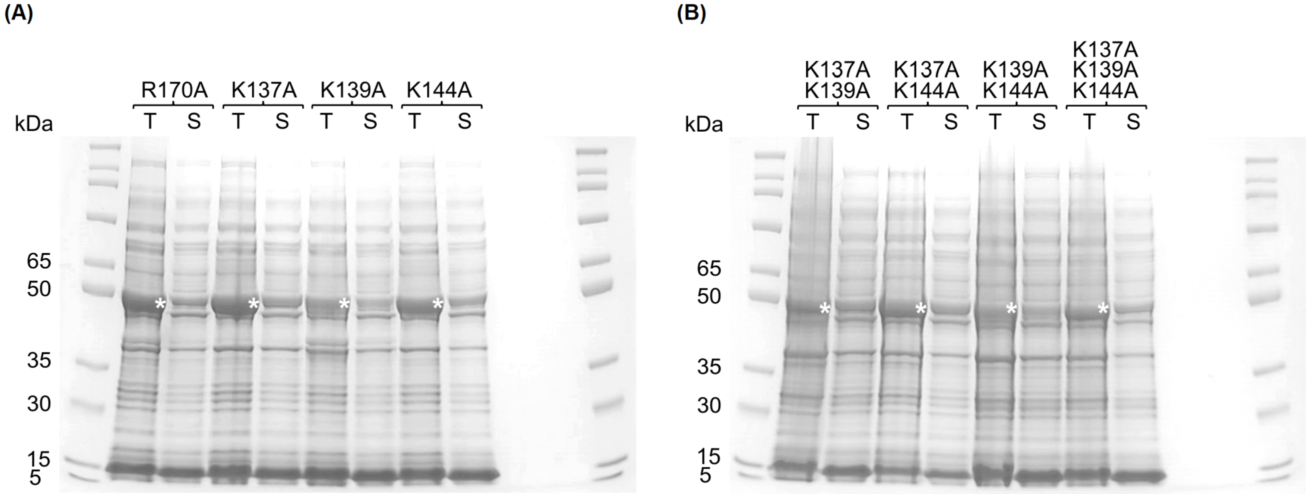


**Supplementary Figure 3.** Image of SDS-PAGE gel showing total (T) enzyme expressed and the corresponding soluble protein (S) of site-directed globupain mutants (A) $R_{170}A$, $K_{137}A$, $K_{139}A$, $K_{144}A$ and (B) $K_{137}A/K_{139}A$, $K_{137}A/K_{144}A$, $K_{139}A/K_{144}A$, $K_{137}A/K_{139}A/K_{144}A$. White asterisks indicate estimated protein size. Protein marker (Broad Multi Color Pre-Stained Protein Standard, Genscript) with indicated molecular weight in kDa are shown to the left and far right in (A) and (B).

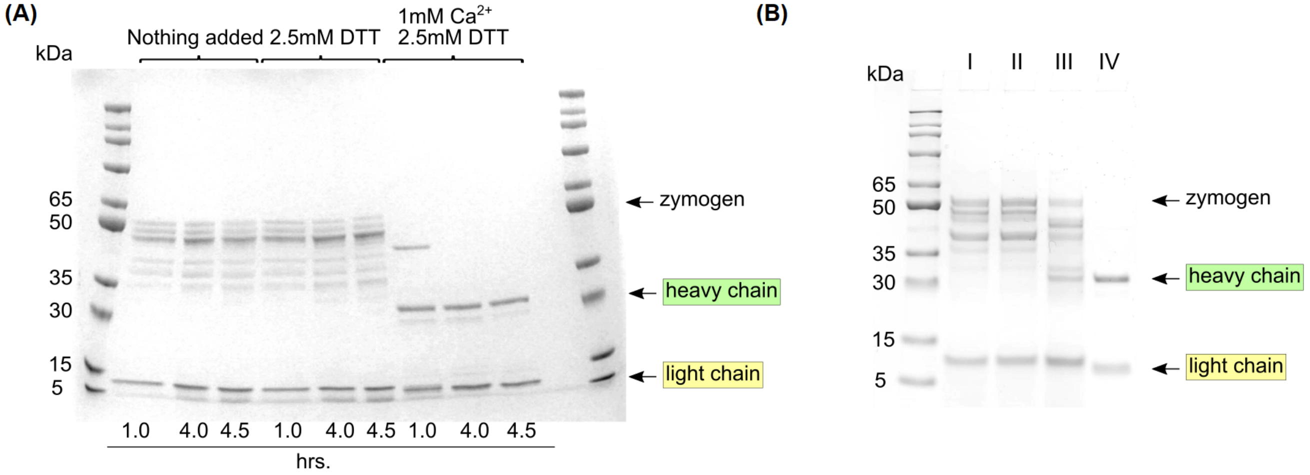


**Supplementary Figure 4.** Effect of calcium and DTT on globupain activation. (A) SDS-PAGE gel showing purified globupain at various time points (1, 4, 4.5h) incubated at 75°C in buffer (20mM tri-sodium citrate dihydrate, 150mM NaCl, pH 5.5). When nothing was added to the buffer, no active heterodimer was observed on the gel. 2.5mM DTT alone did not cleave the zymogen into the active heterodimer but the combination of 1mM $Ca^{2+}$ and 2.5mM DTT fully cleaved the zymogen into the active heterodimer after 4-4.5h. (B) SDS-PAGE gel of purified globupain incubated for 4.5h at 75°C in buffer (20mM tri-sodium citrate dihydrate, 150mM NaCl, pH 5.5) with I. nothing added, II. 1mM EDTA added, III. 1mM EDTA/1mM $Ca^{2+}$/2.5mM DTT added and IV. 1mM $Ca^{2+}$/2.5mM DTT added. Results show that addition of EDTA inhibits the activation of globupain in presence of $Ca^{2+}$ and DTT. Thus, DTT and calcium are required for activation.

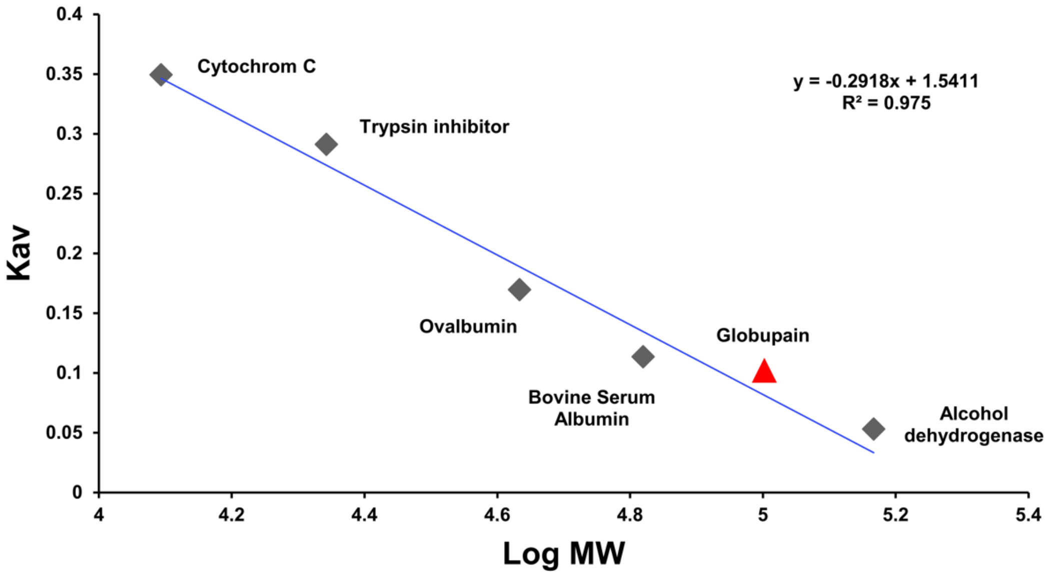


**Supplementary Figure 5.** Determination of oligomeric structure of globupain protein by size-exclusion chromatography (SEC). The SEC analysis was performed using a Superdex 75 10/300 GL prepacked column connected to ÄKTA pure 25 chromatography system (GE Healthcare). The column was equilibrated with a 50 mM potassium phosphate buffer (pH 7.0), 150 mM NaCl and then loaded with a 500 µL sample of globupain protein (1 mg/ml). The flow rate of the run was adjusted to 0.5 mL/min, and the absorbance was measured at 280 nm (mAU, milli-absorbance units). For the experiment, the column was calibrated with proteins of known molecular weight: alcohol dehydrogenase (tetramer), 146,800; bovine serum albumin, 66,000; ovalbumin, 43,000; trypsin inhibitor, 22,000; cytochrome C, 12,400 (Sigma-Aldrich, St. Louis, MO, USA). Dextran blue 2000 (Cytiva) was used to determine the column void volume.

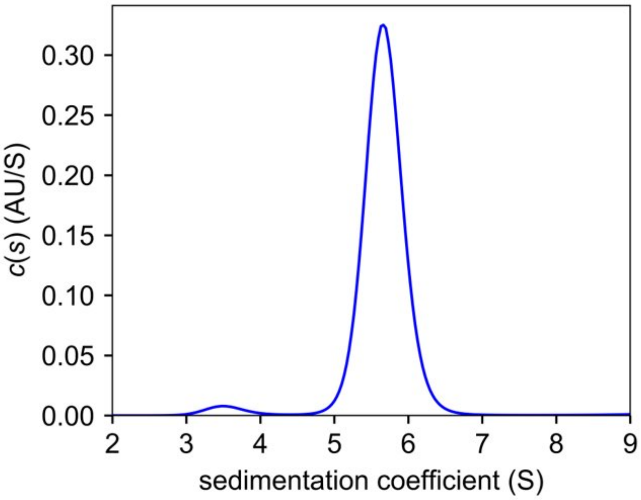


**Supplementary Figure 6.** Sedimentation coefficient distributions of globupain protein as determined by analytical ultracentrifugation. By nonlinear fittings, the average molecular weight was determined as 103,000 for the C11_clostripain sample (sedimentation coefficient 5.67S). This result indicates that the protein exists in solution predominantly (92%) as a dimer. (Experiment parameters: temp. 20 °C, 50 k rpm, scans were collected at 280 nm with 4 min. intervals between scans, proteins partial specific volume V-bar =0,7309 ml/g , buffer density = 1,01395 g/cm3 , buffer viscosity =1,030 mPa·s.)

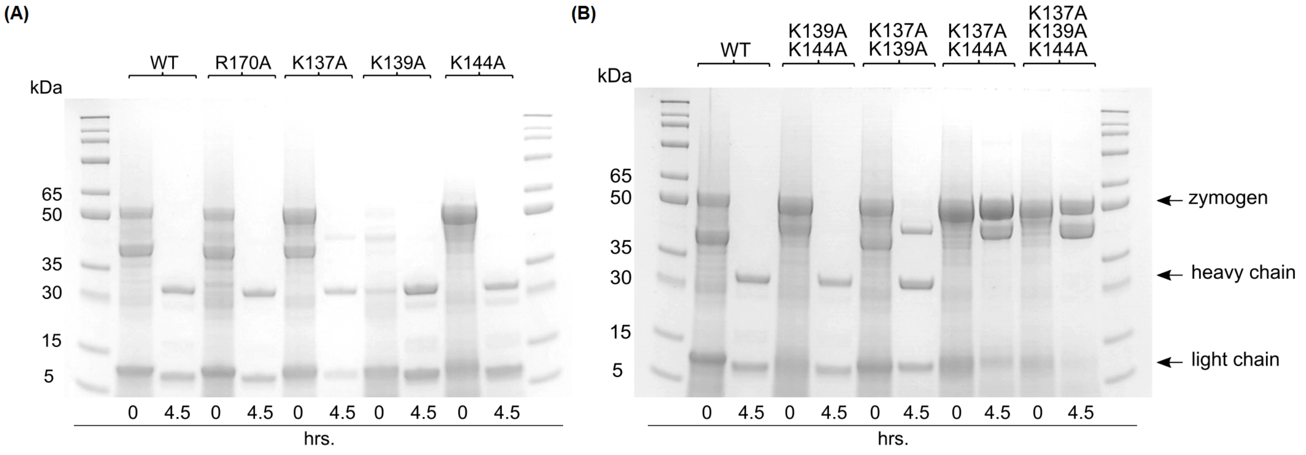


**Supplementary Figure 7.** SDS-PAGE gel image showing enzyme in activation buffer at time 0h and 4.5h incubated at 75°C for (A) WT globupain and site-directed mutants $R_{170}A$, $K_{137}A$, $K_{139}A$, $K_{144}A$ and (B) $K_{137}A/K_{139}A$, $K_{137}A/K_{144}A$, $K_{139}A/K_{144}A$, $K_{137}A/K_{139}A/K_{144}A$. Results show that $K_{137}A/K_{144}A$ and $K_{137}A/K_{139}A/K_{144}A$ failed to produce heavy- and light chain of the active heterodimer.

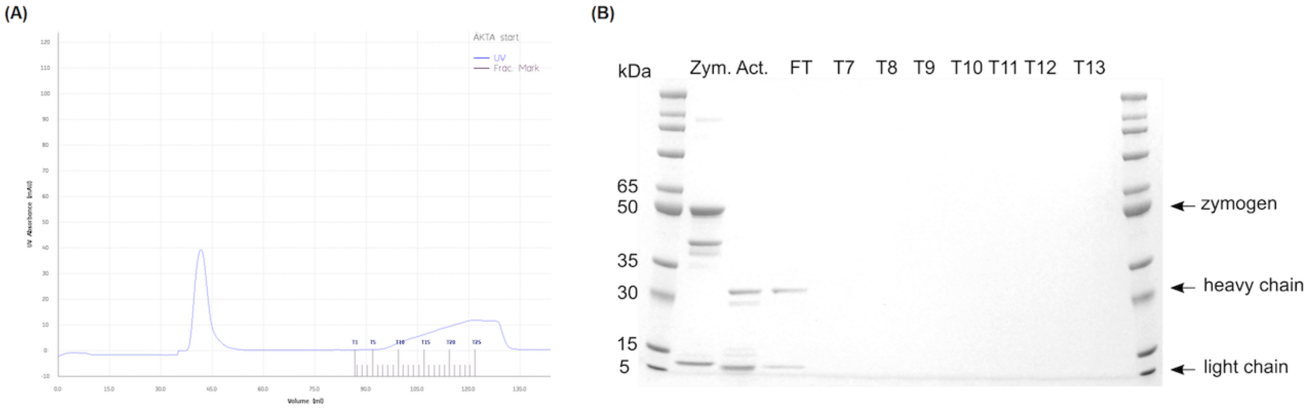


**Supplementary Figure 8.** (A) IMAC purification of activated globupain enzyme showing the UV absorbance read in milli-absorbance unit (mAU) of activated enzyme flowing through the HisTrap HP 5 mL column and failing to elute into the fractions (T1-T25), indicating loss of the 6xHIS-tag. (B) SDS-PAGE gel image of IMAC purification of WT globupain zymogen (Zym.) that was activated (Act.) and loaded onto HisTrap HP 5 mL column showing that the peak at 40mAU in (A) was the activated enzyme (FT). No enzyme was observed in elution fractions T7-T13 with only the imidazole gradient showing in (A) for T5-T25.

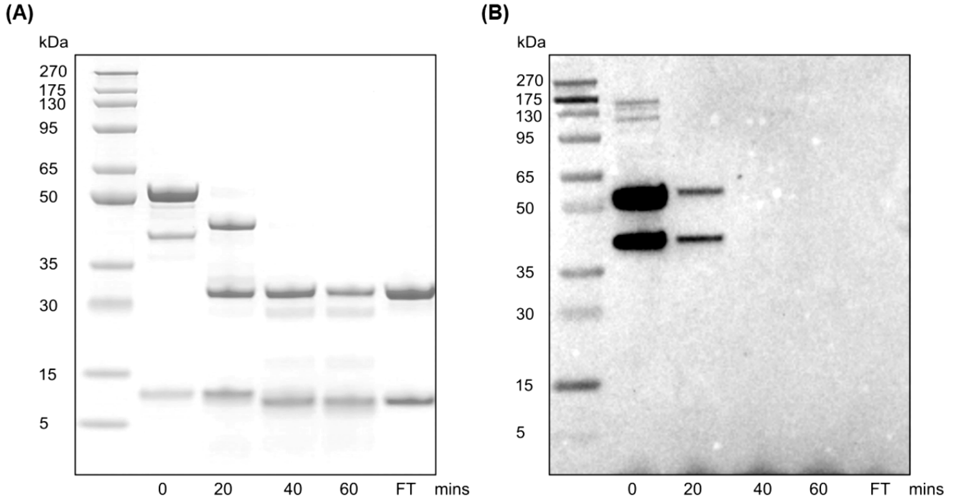


**Supplementary Figure 9.** SDS-PAGE gel of globupain activation time series and Western immunoblotting. (A) Protein bands at 60 minutes of activation conditions at 75°C with 20-minute intervals. Flow through (FT) of IMAC purification showed that the active heterodimer did not bind to the $Ni^{2+}$ affinity column. (B) Western immunoblotting of activation time series with identical setup as in (A). Dark bands indicate the presence of peptides with Histidine-tag. The image shows that the 52kDa globupain zymogen has the His-tag in addition to a 40kDa band which most likely represents an unspecific N-terminal cleave. The active heterodimer does not have the His-tag. Western immunoblotting was performed to validate the observation of activated globupain lacking the C-terminal portion with His-tag. Zymogen and activated globupain heterodimer were transferred from SDS-SurePAGE gel to a nitrocellulose membrane. Blocking was performed by adding 5% fat-free milk dissolved in 50mM Tris buffer, pH 7.5, 150mM NaCl and 0.1% Tween-20 followed by 1h incubation at room temperature. Anti-His-tag mouse monoclonal antibody (OriGene) was used as the primary antibody and incubated overnight at 4°C. The secondary antibody, Anti-mouse IgG Horseraddish Peroxidase linked whole antibody (from sheep) (GE Healthcare) was incubated for 1h at RT. Enhanced chemiluminescence detected by BioRad ChemiDoc™ XRS+ enabled the detection of His-tagged protein treated with SuperSignal West Pico PLUS Chemiluminescent Substrate (Thermo Fisher Scientific).

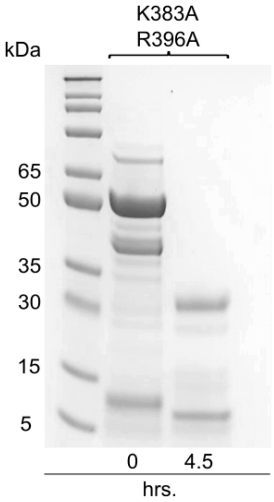

**Supplementary Figure 10.** Image of SDS-PAGE gel of the site-directed mutant $K_{383}A/R_{396}A$ in activation buffer incubated at time 0h and 4.5h at 75°C. The enzyme was able to cut into the active heterodimer.
